# Spontaneous development of cerebrovascular pathology and microinfarcts in a mouse model of sickle cell disease

**DOI:** 10.1101/103309

**Authors:** Hyacinth I. Hyacinth, Cortney L. Sugihara, Thomas L. Spencer, David R. Archer, Andy Y. Shih

**Author notes:** Corresponding Author: Correspondence to: 2015 Uppergate Drive, Atlanta, GA 30322.

## Abstract

Stroke is a dramatic complication of sickle cell disease (SCD) and is associated with aneurysms, moya moya, intravascular thrombi, cerebral hyperemia and increased vessel tortuosity. We show that aged SCD mice spontaneously develop the characteristics features of cerebral vasculopathy seen in human SCD. Thirteen month old Townes SCD mice and age-matched controls had a cranial windows implanted over the somatosensory cortex. Cortical capillaries were imaged using *in vivo* two-photon microscopy after the blood plasma was labeled with a fluorescent dye. Results showed that SCD mice compared to controls, had significantly higher red blood cell (RBC) velocity and capillary vessel diameter. SCD mice also had a significantly higher number of occlusive events in the capillary bed, resulting in more stalling of RBC flow. Microvascular topology was also altered, as SCD mice had significantly higher vessel tortuosity and shorter capillary branch lengths. Finally, *post-mortem* analyses revealed a greater number of cortical microinfarcts, likely caused by vascular occlusion since local tissue hypoxia and blood-brain barrier leakage was prominent. We concluded that aged Townes sickle cell mice spontaneously develop SCD-associated cerebral vasculopathy, and that *in vivo* two-photon imaging is a powerful approach to investigate the mechanisms of vascular complications in SCD.

## INTRODUCTION

In homozygous sickle cell disease, the substitution of valine for glutamic acid in the beta chain of the globin portion of the hemoglobin molecule forms sickle hemoglobin.^1^ Sickle hemoglobin crystallizes during conditions of hypoxemia, leading to defective erythrocytes that are fragile and easily lysed.^2^ The resulting chronic hemolysis and anemia is accompanied not only by exuberant erythropoiesis with increased catabolism and protein synthesis in the individual, ^3^ but also by chronic inflammation secondary to ongoing vascular endothelial injury and recurrent vaso-occlusive events.^4, 5^

Animal models offer valuable opportunities to test therapeutic interventions and management regimes; each model has its own strengths and weaknesses.^6^ Since its introduction in 1997, the Berkeley sickle cell transgenic murine model (S) has served well for the study of some features of human sickle cell disease, especially chronic erythrocyte sickling, tissue hypoxia, hemolytic anemia and extramedullary hematopoiesis.^7^ It has been less well suited, however, for studies of sickle cell chronic lung disease, cerebral vasculopathy, bone marrow infarction or embolization, retinopathy, renal papillary necrosis or splenic atrophy,^8^ due to its genetic background and the non-availability of a humanized control. The histopathological changes of chronic tissue hypoxia and hemolytic anemia in the Berkeley model include infarcts and siderosis in spleen, kidneys and liver; as well as, splenomegaly secondary to exuberant hematopoiesis. Manci et al previously stated that while transgenic sickle cell mice modelled very closely many aspects of human SCD, they failed to show spontaneous cerebrovascular pathology which was well documented as a complication of SCD in humans.^8^ In a recent review, Hillery et al reported that SCD is associated with microvascular cerebral vasculopathy,^9^ which might require time to develop in the SCD mouse model.

Here, we used *in vivo* two-photon laser-scanning microscopy to study cerebral microvascular changes that are associated with age in Townes mice^10^ Previously, we noted that reports of absent spontaneous cerebrovascular changes in sickle cell mouse model were from an experiment utilizing mice between the ages 1 to 6 months.^8^ We therefore hypothesized that cerebral vasculopathy in these mice were dependent upon age. We show that aged (13 months old) Townes mice are a useful model for *in vivo* imaging studies of cerebral vasculopathy and a potential platform for testing therapeutics that may alleviate cerebrovascular disease seen in SCD.

## METHODS

### Animal preparation

We utilized the Townes transgenic mouse model which is a humanized sickle cell (HbSS or SCD) model with a humanized wild-type (HbAA or control). A total of 10 mice, five each of HbAA and HbSS males mice were aged at the Emory pediatrics animal facility and at 13 months of age were transferred from Emory to the Medical University of South Carolina. The mice were allowed to acclimatize for 1 week before surgery and imaging. After acclimatization, the mice were transferred to the lab in pairs for surgery. Each transgenic SCD mouse was processed and imaged alongside an age matched HbAA mouse. We have previously described the methods for preparing and imaging mice using in vivo two-photon microscopy.^11^ However, because of the unique nature of the SCD transgenic mice, we made some modifications to our procedure and thus it bears a short description.

The mice were anesthetized using 2% isoflurane (Patterson Veterinary) in pure oxygen in an induction chamber. They were then transferred to the operating stage where isoflurane was delivered through a nose cone and an adequate depth of anesthesia was maintained via titration by response to a slight pinch of the foot. The skin was cleaned with betadine and then the cranial bone exposed. The periosteum was gently removed using a fine dissecting tools under a dissecting microscope and then dental cement was used to affix a custom-made metal flange for head stabilization during imaging, and to form a head cap to protect the cut skin. A heating pad set at 37°C was used to maintain constant body temperature.

Using a fine drill (Osada) and 0.5 mm drill burr (Fine Science Tools, Foster City, CA), a craniotomy was gently made by removing an approximately 3mm x 3mm piece of skull bone overlying the sensory cortex. The skull bone and subsequently the pial surface was constantly moistened with artificial cerebrospinal fluid avoid desiccation. Agarose gel was placed around the cranial window and a no. 0 glass coverslip laid over the window to overlap slightly the edge of the window. The glass slide was secured with dental cement before imaging was started.

### In vivo imaging procedure

Two-photon imaging was performed with a Sutter Moveable Objective Microscope (MOM) and a Coherent Ultra II Ti:Sapphire laser source. Mice were maintained under light isoflurane (0.5-0.75%) supplied in air (20-22% oxygen and 78% nitrogen, moisturized by bubbling through water) over the duration of imaging. The plasma was labeled by retro-orbital vein injection of 0.03 mL of 2 MDa fluorescein-dextran (FD2000S; Sigma-Aldrich) prepared at a concentration of 5% (w/v) in sterile saline^12, 13^. In a subset of mice, a pulse oximeter (Starr Life Sciences Corp.) was used to ensure that blood oxygen saturation, heart rate and breathing were maintained within normal physiological range with our imaging paradigm. Procedures for blood flow imaging and analysis have been described previously^13^. Wide-field images were collected using a 4X, 0.13 NA objective lens (Olympus UPLFLN 4X) to generate vascular maps of the entire window for navigational purposes. High-resolution imaging of microvessels was performed by using a 20X, 1.0 NA objective lens (Olympus XLUMPLFLN 20XW). High-resolution image stacks of the vasculature were collected across a 326 by 326 µm field and up to a depth of 400-500 µm from the pial surface. Lateral sampling was 0.6 to 0.3 µm per pixel and axial sampling was 1 µm steps between frames.

### Immunohistochemistry

Animals were perfusion fixed with 4% (w/v) paraformaldehyde in PBS. Brain were extracted and post-fixed overnight at 4°C. The entire brain was sectioned because the infarcts were spontaneous and we did not know apriori, the location of infarcts. Brain sections were collected at a thickness of 50 µm and stored in PBS with 0.2% sodium azide. Anti-NeuN primary antibody from mouse host (MAB377; Millipore, Billerica, MA) was diluted in buffer consisting of 10 % (v/v) goat serum (Vector Labs), 2 % (v/v) Triton X-100, and 0.2 % (w/v) sodium azide. Free-floating sections were then incubated overnight under slow nutation at room temperature. The following day, sections were washed in 50 mL of PBS for 30 minutes on an orbital shaker, incubated with Anti-mouse Alexa 594 secondary antibody for 2 hours, washed again in PBS, mounted and dried on slides for 30 minutes. All slides were then sealed with Fluoromount-G (Southern Biotechnology Associates Inc.) and a No. 1 glass coverslip (Corning). Staining for other targets such as CD31 (550274 BD Biosciences) for the endothelium used a similar protocol as above. For pimonidazole hydrochloride (HP3-100Kit, Hypoxyprobe^TM^; Hypoxyprobe.com) immunostaining of hypoxic tissue^14^, pimonidazole was injected via the retro-orbital route one hour prior to perfusion fixation at a concentration of 60 mg/kg in a volume of 100 µL saline. The bound hypoxyprobe adduct is detected using rabbit anti-hypoxyprobe antibody. Finally cell nuclei were labeled by adding Hoescht 33342 (Sigma-Aldrich) at a ratio of 1:100 to the tissue in secondary antibody solution, 10 - 30 mins prior to washing.

### Microvessel Analysis

#### Vessel masking

161×161×100 µm image volumes, collected with two-photon microscopy, were processed into binary masks using ImageJ software. Separate masks were created for skeletonization and vessel volume analysis, respectively. Processing of images occurred in the following order for each analysis:

#### Vascular Volume Mask

(1) Image converted to 8-bit. (2) 3D smoothing of 8-bit image utilizing a Gaussian filter with sigma set to 1.0. (3) Thresholding of image to include all pixels of between the intensity values 15 to 355. (4) Creation of binary mask based on the thresholded image. (5) 3D erosion algorithm with an isovalue of 155, applied three times to more accurately reflect the space occupied by the intraluminal dye. Using custom MATLAB software, the volume occupied by voxels with a value of 1 was divided by the volume of all voxels examined, *i.e.*, 0 and 1. The calculated value may under-estimate the total intravascular volume, since blood cells are not labeled by the FITC-dextran dye.

#### Microvessel skeletonization

(1) Image converted to 8-bit. (2) 3D smoothing of image utilizing a Gaussian filter with sigma set to 1.0. (3) Thresholding of image to include all pixels of between the intensity values of 15 to 355. (4) Creation of binary mask based on the thresholded image. (5) A 5×5×5 µm, 3D median filter applied to the binary mask to eliminate noise. (6) 3D skeletonization algorithm run on filtered binary mask. “Skeletonize (3D)” (http://fiji.sc/Skeletonize3D) and “Analyze Skeleton” (http://fiji.sc/AnalyzeSkeleton) plugins for Fiji software used for these procedures. (7) Vessel segments with branch length less than 5 μm, the average diameter of a capillary, was removed to eliminate branches fabricated by the skeletonization algorithm when vessels of larger diameter, *i.e.* > 10 μm, were present in the image stack. (8) Using custom MATLAB and BoneJ plugin, the microvascular branch volume, length, and Euclidean distance between the tips of branch points were determine.

#### Quantification of vasodynamic and cellular parameters

User-defined line scans,^15^ videos and Z-stacks obtained from each mouse from each genotype group were analyzed. Using custom MATLAB codes and Fiji/Image J software, an unbiased biased evaluator analyzed both the line scans and Z-stacks. Z-stacks were analyzed for the proportion of vessels with intravascular occlusive events (defined *a priori* as an unlabeled intravascular object that is 10× or more the diameter of an RBC; **Supplemental Figure 1a**), proportion of vessels with blood flow stalls (defined as a total cessation of blood flow on z-stacks or videos) and number of RBCs arrested (defined as an RBC that is stationary for 1 or more seconds; **Supplemental Figure 1b**) per unit time and unit length of vessel segment). Further, line scans were analyzed for RBC velocity using the custom MATLAB codes.^16^ Capillary diameter was measured manually (by eye) using ImageJ software. Red blood cell flux was calculated using the formula^14^:

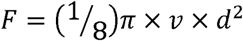

Where *F* is RBC flux or cortical blood flow through a vessel in nL/sec, *v* is velocity of flow and *d* is the diameter of the vessel. Although a technical limitation of the above formula is that it assumes laminar flow which is not the case in capillaries which has a “single file flow”, but nevertheless allows an estimation of the impact of lumen diameter changes co-occurring with RBC velocity change. Euclidean distance, a measure of vessel tortuosity and the average microvascular branch length were obtained from the analysis of skeletonized 3D image reconstruction of the z-stacks.

### Statistics Analysis and image processing

Data analysis for comparison between SCD mice and controls were done using the t-test function of GraphPad Prism software (GraphPad Software Inc, La Jolla, CA). Image processing, analysis and scaling, and video processing was done using the combination of Adobe CS6 (Adobe Systems Co., San Jose, CA), Fiji/Image J (<u>http://jenkins.imagej.net/job/Stable-Fiji/lastSuccessfulBuild/artifact/fiji-nojre.zip</u>) and Microsoft Movie Maker (Microsoft® Corp. Redmond, WA). Quantitative results are presented using bar graphs comparing sickle to control mice and with a p-value of <0.05 considered statistically significant. Qualitative data are presented as representative videos or an array of histochemical images.

## RESULTS

### Cerebrovascular morphology and cerebral vasodynamics in SCD mice

We used 13-month old male Townes control and SCD mice in this study. Cerebral vasodynamic measurements were obtained by *in vivo* two-photon blood flow imaging, using methods previously described^13, 15^ Analysis of cerebral vasodynamic measurements revealed that sickle cell mice had significantly higher capillary RBC velocity (0.73 mm/sec vs. 0.55 mm/sec, p = 0.013) compared to controls (Fig. 1A). In addition, SCD mice also had significantly larger capillary vessel diameter (4.84 µM vs. 4.50 µM, p = 0.014) compared to controls (Fig. 1B). Since RBC volume flux is a function of vessel diameter and RBC velocity, sickle cell mice had significantly higher RBC volume flux (0.015 nL/sec vs. 0.010 nL/sec, p = 0.021) compared to controls (Fig. 1C).

**Figure 1.**
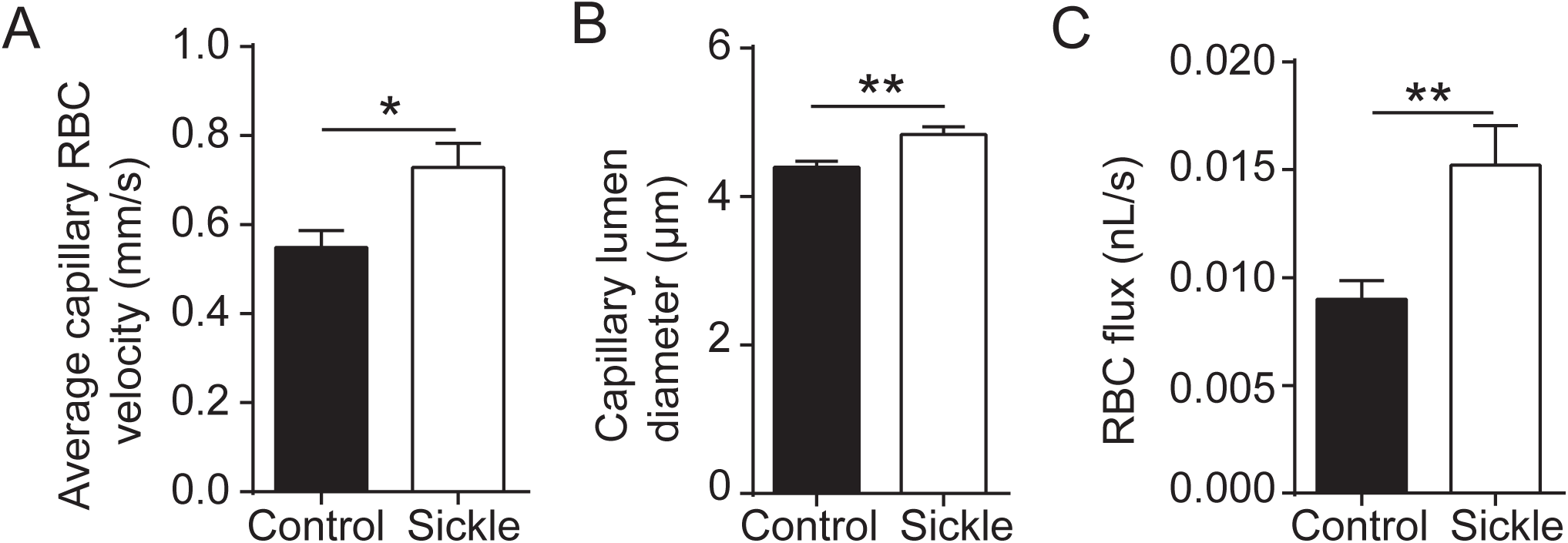
Altered capillary vasodynamics in SCD mice. Red blood cell velocity (A), mean capillary diameter (B), and red blood cell flux (C) was significantly higher among SCD mice compared with controls (p = 0.013, 0.014, 0.021 for RBC velocity, capillary diameter and flux, respectively; t-test). Data is presented as mean ± s.e.m.

Sickle cell mice also had gross evidence of altered cerebrovascular topology (**Supplemental videos 1a-d**). Specifically, we determine the Euclidean distance of capillary branch-points, *i.e.*, the 3D distance between two branch-point vertices, which has been shown to indicate the level of tortuosity of a vessel.^17, 18^ Euclidean distance was significantly shorter in sickle cell mice (21.7µm vs. 38.3µm, p<0.0001) compared to controls (Fig. 2A), indicating that the SCD mice had significantly more tortuous capillary network. Consistent with this, the average length of capillaries between branch-points was also significantly shorter in SCD (15.74µm vs. 51.24µm, p = 0.0065) compared to control mice.

**Figure 2.**
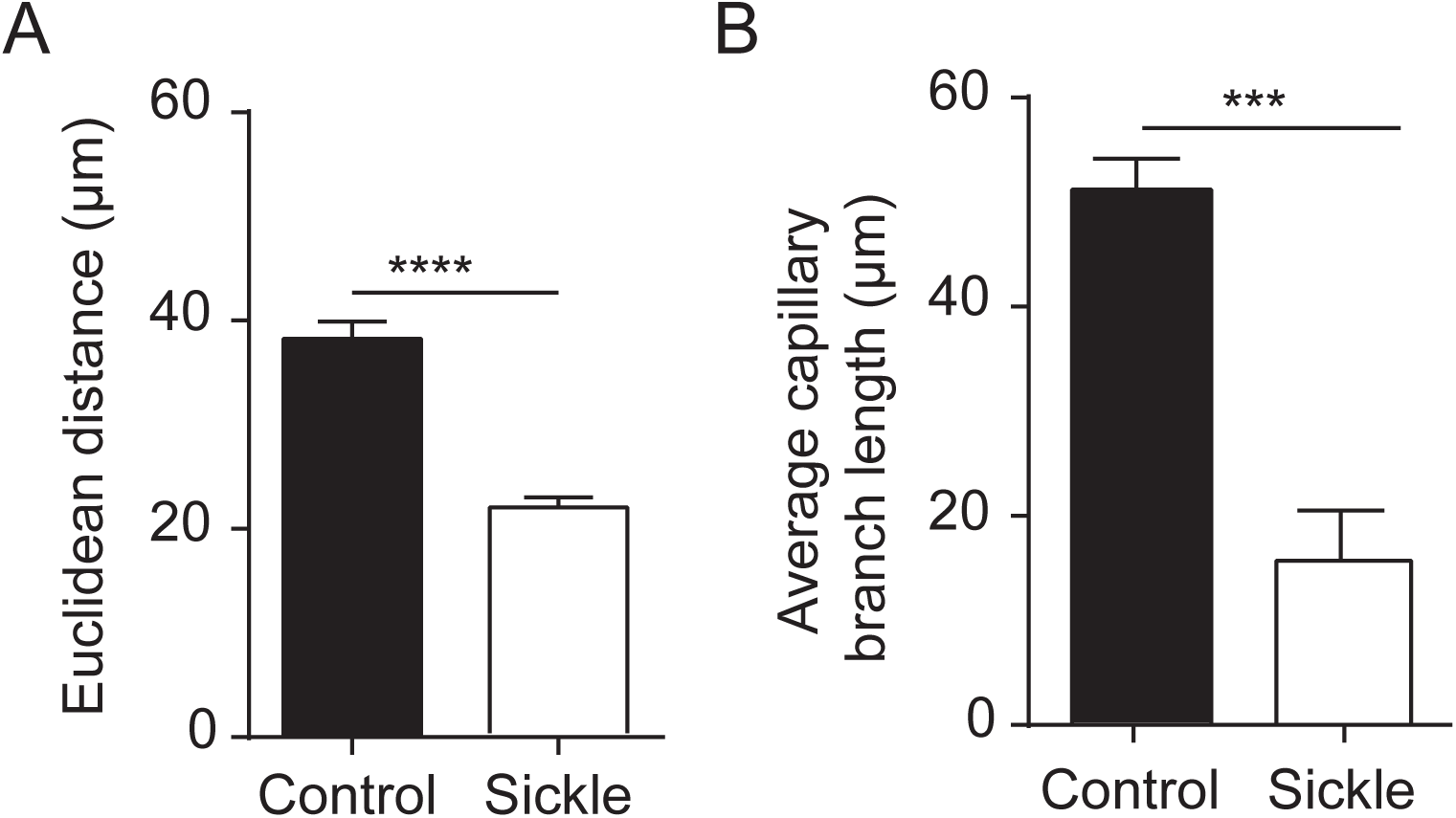
Abnormal capillary topology in SCD mice. (A) The Euclidean distance between capillary branch points, an indication of the degree vessel tortuosity, is significantly shorter in SCD mice (p<0.0001; t-test). (B) SCD mice also have significantly shorter capillary branches compared to controls (p = 0.001; t-test).

**Figure 3.**
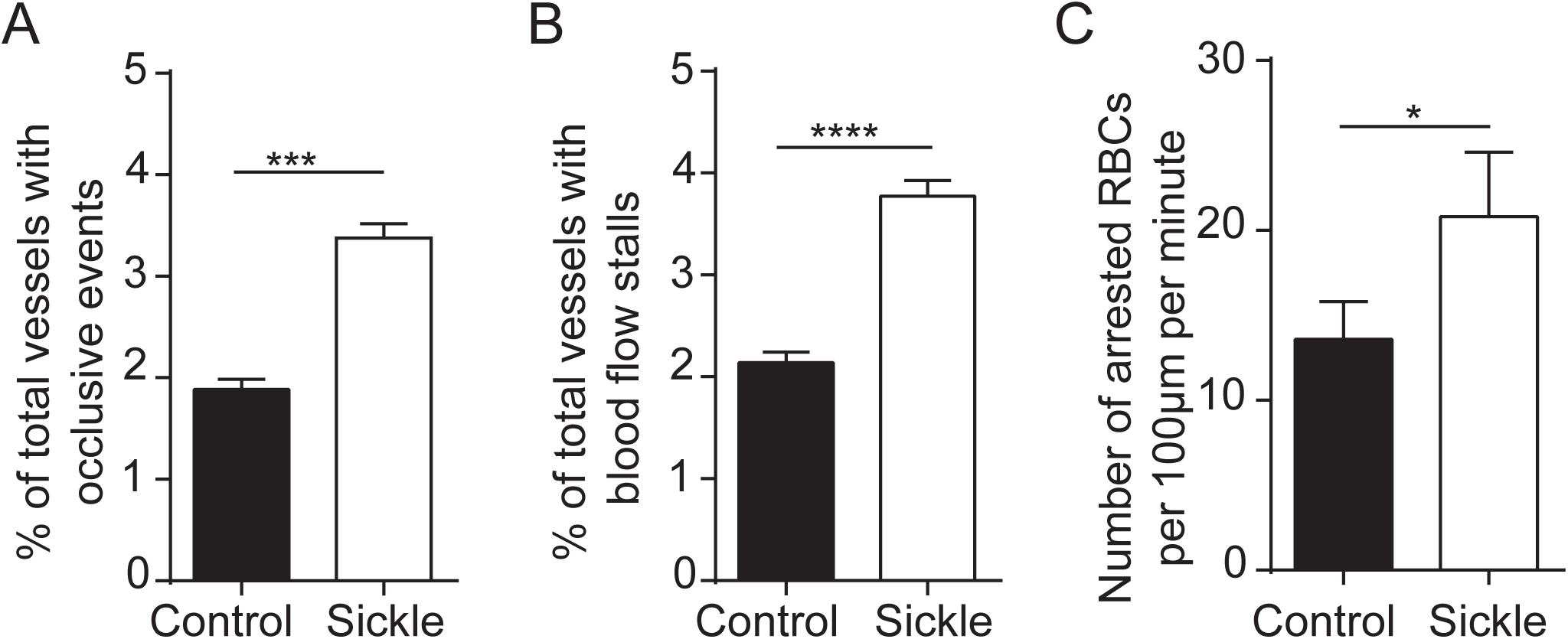
Evidence of cerebral vasculopathy in SCD mice. (A) Higher number of occlusive events (A) blood flow stalls (B) and arrested RBCs (C) in SCD mice, compared to control mice.

### Sickle cell mice compared to control developed spontaneous intravascular pathologies

Using images from line scans and also 3D image stacks, an assessor blinded to the genotype of the mice counted the number of vessels with occlusive events (clots or adherent cells that are ≥ 10× the size of an RBC; **supplemental video 3a** for SCD, **supplemental video 3b** for control) or RBC stalls (capillary segments with stationary RBC). These measurements were then expressed as a percentage of the total number of vessels examined. SCD mice had a significantly higher percentage of cerebral vessels with occlusive events (3.4% vs. 1.9%, p <0.0001) compared to controls. Similarly, sickle cell mice had significantly higher percentage of cerebral vessels with RBC flow stalls (3.8% vs. 2.1%, p < 0.0001) compared to controls. Consistent with the higher incidence of flow disturbances in SCD mice, there was a significantly greater number of arrested RBCs (20.8 vs. 13.6, p = 0.036) per 100µm of vessel length per minute, among SCD mice compared to controls. An arrested RBC was defined as one that remained motionless for 1 second or longer. Thus, SCD mice exhibited greater cortical cerebral vasculopathy compared to controls.

### Spontaneous development of cortical microinfarcts in sickle cell mice

After completion of two-photon imaging, mice were injected intravenously with Hypoxyprobe, a marker that binds tissues with less than 10 mmHg pO_2_ content, providing an immunohistochemical target to identify hypoxic brain regions. Brains were then extracted, sectioned and immunolabeled for Hypoxyprobe, along with markers of neuronal viability, NeuN, and endothelial integrity, CD31. The occurrence of cortical microinfarcts was a prominent feature in the brains of SCD mice (Fig. 4). Microinfarcts manifested as localized regions devoid of NeuN immunoreactivity, indicating loss of neuronal viability (Fig. 4a-d; right panels). They closely resembled the column-like shape of microinfarcts induced by direct occlusion of cortical penetrating arterioles in mouse brain.^19^ Furthermore, consistent with the idea that spontaneous microinfarcts arise from the blockage of cortical penetrating arterioles, we observed changes consistent with a tissue response to ischemia. This included: *i)* local extravasation of the intravenous fluorescent dye, FITC-dextran, confirming blood-brain barrier leakage (Fig. 4a, b, e; right panels), *ii)* increased binding of Hypoxyprobe in the microinfarct core (Fig. 4c, d; right panels), and *iii)* local degeneration of the vascular endothelium (Fig. 4e; left panel). We assessed the frequency of microinfarcts in each group by examining all brain sections from SCD (N = 8) and control (N = 7) mice, encompassing ~100 sections per mouse spanning the entire cerebrum. This revealed a 2.5-fold higher frequency of microinfarcts in the SCD brain versus the brains of control mice.

**Figure 4.**
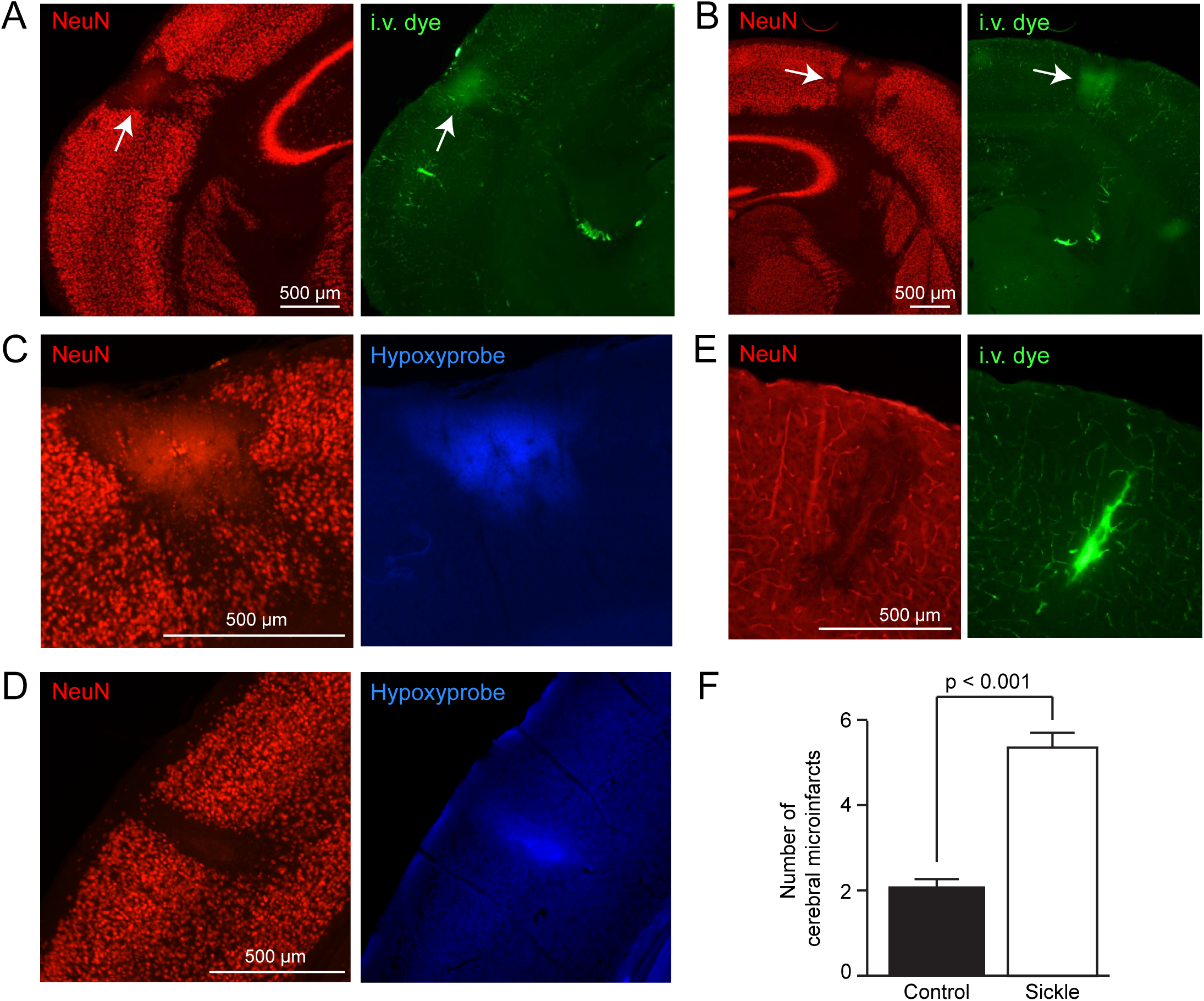
Spontaneous occurrence of cerebral microinfarcts in SCD mice. (A, B) Representative images of cortical microinfarcts in SCD mice. Regions devoid of NeuN staining, a marker of neuronal viability, mark the boundaries of the microinfarcts. Extravasation of the intravenous dye, FITC-dextran, in and around the microinfarct core is consistent with blood-brain barrier disruption. (C, D) Hypoxyprobe labeling marks regions of tissue hypoxia. (E) Reduced CD31 staining indicates endothelial damage in the microinfarct core. (F) SCD mice had a 2.5 fold higher frequency for microinfarcts compared to controls.

## DISCUSSION

In this study, we show that aged Townes sickle cell mice spontaneously develop cerebrovascular pathologies that are akin to those seen in human sickle cell disease. The observance of cerebral vasculopathy in children with sickle cell disease prompted prior efforts to study the cerebrovasculature of sickle cell mice that were 6 months old or younger.^8^ However, cerebrovascular disease in general is an age-related phenomenon, and thus the goal of this study was to examine cerebrovascular function in aged SCD mice.

We observed a significantly higher RBC flow velocity in sickle cell mice compared to controls, which recapitulates what had been demonstrated in children with sickle cell disease.^20^ A high RBC velocity is associated with shortened RBC transit time, which could negatively impact oxygen extraction, with significant neurological consequences as is the case with SCD.^20-23^ In addition, we noted that SCD mice have greater vascular tortuosity and significantly shortened capillary branches. This structural change may contribute to the greater number of RBC stalls we observed in the capillary beds of SCD mice. This finding also supports earlier reported observations of ineffective collateral vessel formation, also known as moya moya.^24^ Moya moya is associated with a significantly increased risk for stroke among individuals with sickle cell disease.^25^

One key observation was that SCD mice had a 2.5-fold increased frequency for cerebral microinfarcts compared with controls, as well as an increased incidence of intravascular occlusive events observed *in vivo*. The significance of this finding is still unknown since a similar observation have not been reported in children or adults with sickle cell disease. However, in the context of vascular cognitive impairment, atherosclerosis or atrial fibrillations can lead to the generation of microthrombi that then circulate into the brain to occlude cerebral small vessels. (PMID: 27539302) ^26^ This is thought to be a source of microinfarcts in the aging brain, and may also play a role in development of microinfarcts during SCD. Since microinfarcts have recently been linked to increased risk for dementia,^27^ it is possible that the accumulation of these lesions may also contribute to cognitive decline in SCD.^28-30^ Our discussion on these microinfarcts is limited by the fact that we did not ascertain their origin. However, their remarkable resemblance to microinfarcts induced by the occlusion of single penetrating arterioles by photothrombosis, suggests that they arise from occlusion of penetrating arterioles. Further, the detection of local hypoxic tissue further suggests that microinfarct in SCD mice are a consequence of vascular occlusion.

In conclusion, we have shown the utility of a new approach to study the cerebrovascular consequences of sickle cell disease, using combined in vivo two-photon laser microscopy with post-mortem immunohistochemistry in an aged sickle cell mouse model. Using this approach, we have been able to measure abnormalities in microvascular blood flow structure associated with sickle cell disease with high precision. This model system has potential for the longitudinal study of structural and molecular events involved in the evolution of SCD associated cerebrovascular diseases.

## Acknowledgements

### Author Contribution

HIH and AYS designed the experiment, HIH, CLS, TLS, DRA and AYS performed experiments and data analysis, HIH, CLS, TLS, DRA and AYS wrote the manuscript. HIH and AYS performed final critical review. All authors endorsed the submission of this manuscript.

### Acknowledgement

This study was partially funded by Emory University Pediatrics Pilot Grant (HeRO to HIH) and NIH/NHLBI – U01HL117721. Also grants to A.Y.S. from the NINDS (NS085402), the Dana Foundation, South Carolina Clinical and Translational Institute (UL1TR000062), and an Institutional Development Award (IDeA) from the NIGMS under grant number P20GM12345.

